# Evolutionary origin of disease and complexity: A nonequilibrium thermodynamic solution

**DOI:** 10.1101/356493

**Authors:** Lauren Gerard Koch, Steven L. Britton

**Affiliations:** Department of Physiology & Pharmacology, The University of Toledo, Toledo, OH; Department of Anesthesiology, University of Michigan, Ann Arbor, MI; Department of Molecular and Integrative Physiology, University of Michigan, Ann Arbor, MI

## Abstract

Our long-term goal was to develop realistic animal models of complex disease that are underwritten by fundamental principles. Along this path we noted a literature demonstrating that **low exercise capacity is a stronger predictor of death relative to all other clinical conditions including diabetes and smoking.** From this linkage we formulated the Energy Transfer Hypothesis (ETH): *Variation in capacity for energy transfer is the central mechanistic determinant of the divide between disease and health.* As an unbiased test of the ETH we reasoned that: divergent artificial selection of rats based on low and high intrinsic treadmill running capacity would yield contrasting models of capacity for energy transfer that also divide for disease risks. Thirty-five generations of selection produced Low Capacity Runners (LCR) and High Capacity Runners (HCR) that differ in running capacity by over 8-fold. Selection failed to disprove the ETH: disease risks segregate in the LCR, and the HCR demonstrate resistance to disease. For mechanistic explanation of the ETH we postulate that evolution, life, and thus disease follow the same entropic path of the other atoms and molecules of the universe. That is, 1) all systems tend towards Maximal Entropy Production, 2) entropy can temporarily decrease locally and form order such as life, 3) life developed with a genome as a path for coding for replication of energy metabolism units, and 4) evolution selects, in a constantly changing environment, for features that enhance life’s capacity for energy transfer. That is, Maximal Entropy Production is the driving force mediating evolution. Thus, for each species, consider that there is phenotypic variation for this capacity that underlies the divide between health and disease. One goal of this paper is to stimulate new cross-disciplinary tests of the ETH of disease.

## Statement of the problem

In 1986 Koch and Britton speculated that commonly used animal models of complex disease were too simplistic to enable progress in understanding conditions such as diabetes, heart failure, or cancer. Our goal was to devise a useful animal model system to understand the divide between disease and health. As a heuristic we formulated four simplistic “rules” for the minimal features of an animal model of complex disease. We thought a useful model should: 1) harbor clinically relevant phenotype(s), 2) be polygenic and thus heritable, 3) be modulable by positive (exercise) and negative (high fat diet) health environments, and 4) be explained by fundamental scientific principles (1). From our new “rules-based” perspective we considered that the study of widely used animal models of disease might actually be moving us further from useful knowledge because of their artificiality. Widely used overly simplistic examples include: streptozotocin for diabetes mellitus, acute coronary artery ligation for heart failure, gene knock in/knock out to reveal “disease-related” phenotypes and, ethylnitrosourea (ENU) mutagenesis with screening for mutants with clinically-relevant phenotypes. It was clear to us that these approaches did not emulate the polygenic nature of complex diseases and that alternative paths were needed.

## Formulation of a hypothesis

We had noted the emergence of a strong statistical association between low capacity for energy transfer and high risk for disease in the clinical literature that started about 30 years ago (2, 3) and that is now extensively confirmed in large-scale contemporary studies (4, 5). Indeed, it is widely accepted that low capacity for energy transfer is a larger risk factor for death relative to all other clinical conditions including diabetes, smoking, and coronary artery disease (5). From this linkage of complex disease risks with low capacity for energy transfer, we initiated the *Energy Transfer Hypothesis (ETH): Variation in capacity for energy transfer is the central mechanistic determinant of the divide between disease and health* (1). Our long-term association with quantitative geneticist John P. Rapp (6) led us to consider that artificial selective breeding could possibly be used as a tool for first test of the ETH. That is, as an unbiased test of the ETH we reasoned that: *divergent (two-way) selection of rats based on low and high intrinsic treadmill running capacity would yield contrasting models of capacity for energy transfer that also divide for disease risks.*

The strong clustering of morbidities with low physical fitness suggests common causality. While it is widely hypothesized that some aspect of mitochondrial dysfunction is the mediator of the disease-fitness connection, a general mechanistic explanation for this association has not been defined (7). From the beginning we realized the risk of addressing the writ large nature of the Energy Transfer Hypothesis and using selection as a tool. We could not tell if we were closer to Karl Popper (“a hypothesis must make predictions to be falsifiable”) or Wolfgang Pauli (the hypothesis is “not even wrong”) (https://en.wikipedia.org/wiki/Notevenwrong).

## Long-term selection for energy transfer capacity

By 1996 we had gathered enough resources to start selection for low and high intrinsic (i.e., untrained) running capacity. We consider this intrinsic because the rats were selected based upon running tests on each of just five consecutive days when 12 weeks old. The founder population was 168 N/NIH genetically heterogeneous rats provided by Carl Hansen and Karen Spuhler (8). Selection for low and high capacity was based upon distance run to exhaustion on a motorized treadmill using a velocity-ramped running protocol. The starting velocity was 10 m/min and was increased by 1 m/min every 2 min (slope was constant at 15°). We considered this protocol the “rat equivalent” of the Bruce Protocol used clinically (9). At each generation, within-family selection was practiced using 13 families for both the low and high lines. A rotational breeding paradigm maintained the coefficient of inbreeding at less than 1% per generation(10)

As expected, because we started with a genetically heterogeneous population (11), the rats responded to artificial selection. On average by generation 35 of selection (18 years) the lines differed by greater than 8fold for intrinsic maximal running distance (Figure 1). Selection progressed successfully for both lines; formal regression analysis of distance over generations yielded a *p < 2.2×10*^*−16*^ (i.e., at the machine epsilon in language r) in support of a non-zero slope for both lines. These results are reminiscent of the sustained responsiveness seen in other selectively bred lines. For example, selection for oil concentration in maize continues to increase for over 100 generations (12). The long-term response to selection suggests that the running trait is influenced by many interacting genes.

**Figure 1.**
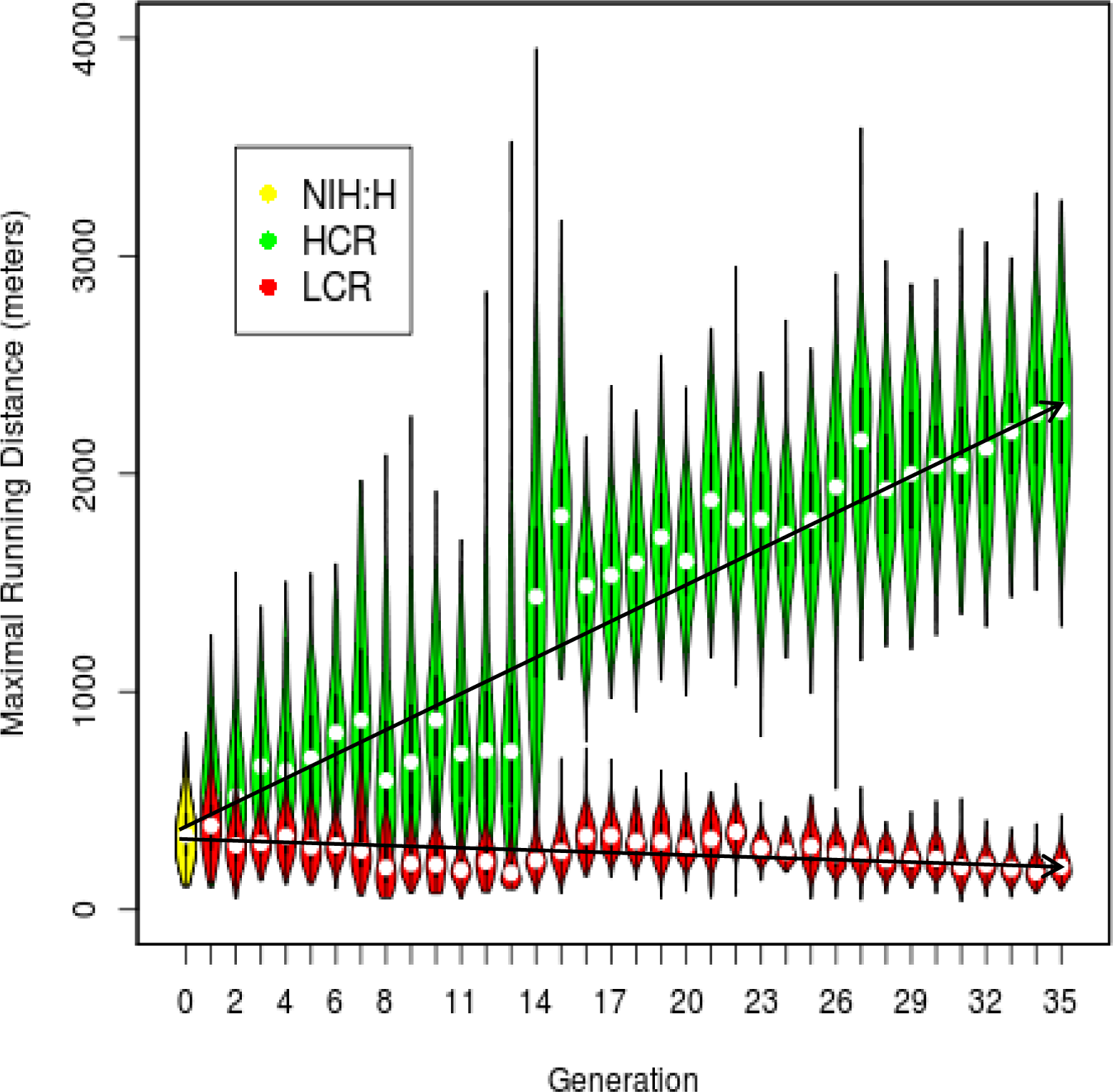
Response to selection for 35 generations. Each symbol represents the distribution of running distance for each generation in each line. The symbols for each generation combine box plots and kernel density plots to depict the observed probability density. Males and females combined (n =13,886 rats). NIH:H are the founder stock rats (31).

## Selection did not falsify the ETH

Disease risks segregated with selection for low capacity exercise and resistance to risks segregated with selection for high capacity exercise. Table 1 shows 12 of the major complex disease risks the LCR harbor relative to the HCR including advanced aging, diminished longevity, metabolic syndrome, and increased susceptibility to ventricular fibrillation. For environment, the LCR are quite susceptible to weight gain on a high fat diet and the HCR are resistant (13).

**Table 1.** Disease Risks (LCR relative to HCR)

| **Table 1. Disease Risks (LCR relative to HCR)** |  |  |
| --- | --- | --- |
| Susceptibility to cancer (19) | Fatty liver disease (20) | Low physical activity (21) |
| Decreased OXPHOS (22) | Alzheimer's neurodegeneration (23) | ↓ Hippocampal neurogenesis (24) |
| Metabolic syndrome (25) | ↑ Susceptibility to cerebral hemorrhage (26) | Mechanism of aging: DNAPk (27) |
| Disordered sleep (28) | ↑ Vulnerability to ventricular fibrillation (29) | Premature aging (30) |

## A theoretical base

In 1966 Francis Crick declared that: “The ultimate aim of the modern movement in biology is to explain all biology in terms of physics and chemistry”.

We took Crick’s broad statement as motivation to originate an approach to understanding complex diseases that is: 1) a deeper extension of conventional molecular biologic reductionism, and 2) based upon principles from non-equilibrium thermodynamics and evolution. The central theme is that evolution, life, and thus disease follow the same entropic path of the other atoms and molecules of the universe.

We sought a path for explanation of the ETH at the mechanistic level. For this, we applied fundamental properties of matter by integrating ideas from Robert Endres (14), Hans Krebs (15), and Peter Mitchell (16). The ideas can be summarized in four statements: 1) Net entropy (S) is always increasing and systems tend towards Maximal Entropy Production (MEP). 2) Entropy can temporarily decrease locally and ordered systems can form (order from disorder) and these ordered systems can dissipate energy faster than non-ordered systems. 3) Among the many possible ordered systems, what we term life developed with a genome as a path for coding for replication of energy metabolism units. 4) Evolution selects, in a constantly changing environment, for features that enhance life’s capacity for energy transfer. Thus, for each species, consider that there is phenotypic variation for this capacity that underlies the divide between health and disease (Figure 2). Our artificial selection outcome is consistent with these arguments.

**Figure 2.**
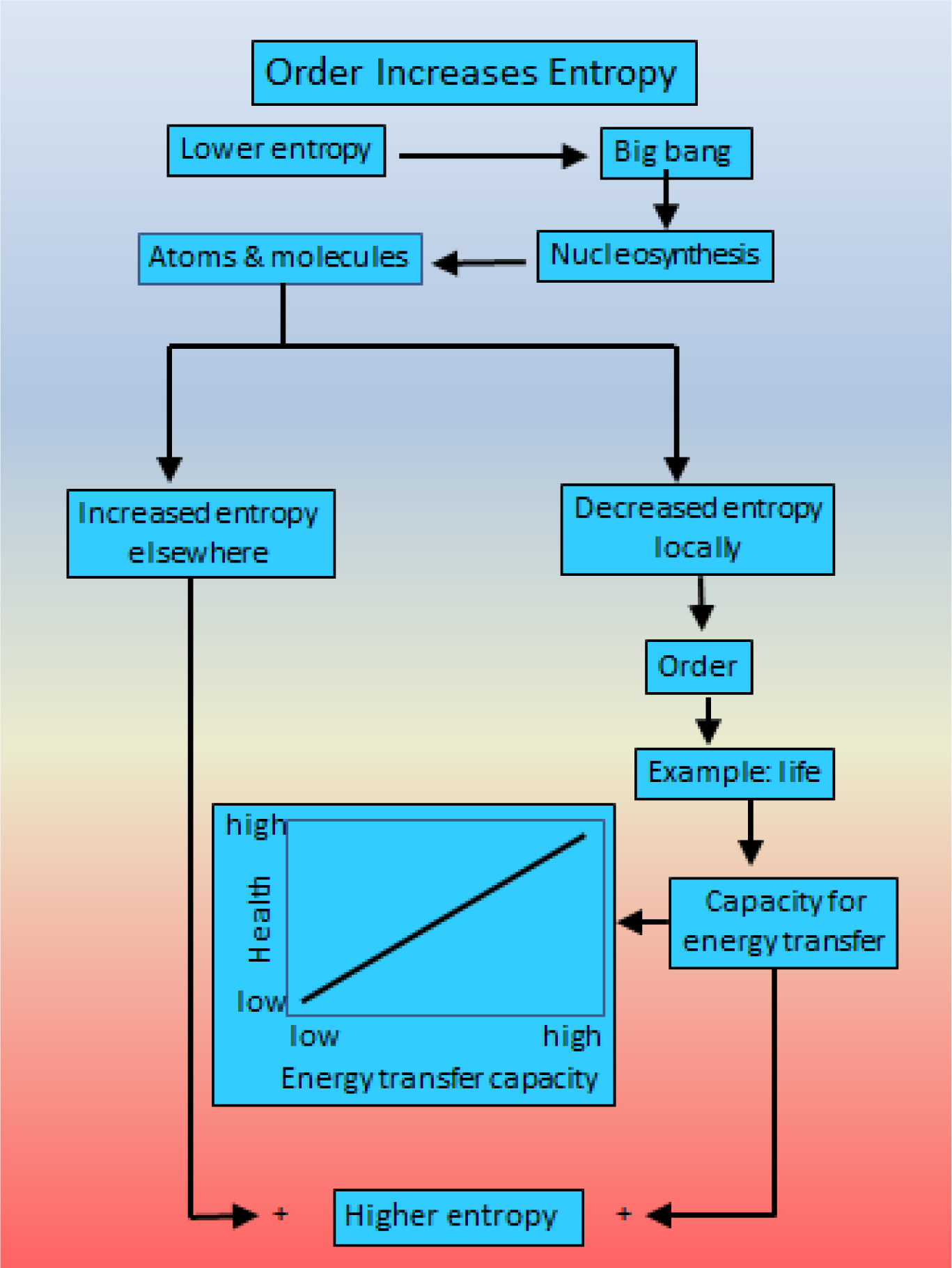
Time zero for biology can be considered as equivalent to the Big Bang and the subsequent formation of atoms (nucleosynthesis) that displayed the property of Maximal Entropy Production (MEP). Locally entropy can decrease (at the expense of increased entropy elsewhere) and ordered systems such as life can form. We postulate that attainment of MEP is the driving force underlying evolution. Thus, for each species, consider that there is phenotypic variation for this capacity that underlies mechanistically the association of low health with low capacity for energy transfer.

## An Extension and meshing of new ideas

Jonathon Pritchard and colleagues (17) re-evaluated the genetic contribution to complex traits and hypothesized that there is an extremely large number of causal variants with tiny (non-zero) effects on height and other complex traits. Moreover, these variants are spread widely across the genome, such that most 100-kb windows contribute to variance in height. They refer to this hypothesis as an “omnigenic model” and estimate that more than 100,000 SNPs exert independent causal effects on height. This type of function is consistent with our above theoretical arguments. That is, the emergence of life represented pathways for enhanced energy transfer that utilized the entire genome for moment-to-moment infinitesimal adjustments to environment. Pritchard proposes that disease risk is driven by genes with no obvious connection to disease that is propagated through regulatory networks to a much smaller number of core genes with direct effects. In refinement, we propose that disease risk is driven by genes that mediate energy transfer capacity that is concealed by the miniscule scale of each accumulative genetic change that occurs with evolution.

## Addendum

The central concept of natural selection is the evolutionary fitness of an organism. Fitness is an organism’s ability to survive and reproduce, which determines the size of its genetic contribution to the next generation. In a large reductionist step we propose that operation of Maximal Entropy Production is the driving force underlying evolution in the sense that it provides the energy for survival and reproductive functions. Of course, at the next level down we do not have specific explanation for the operation of Maximal Entropy Production as is also true for the quantum properties of teleportation, superposition, and entanglement (18).

We trust this succinct perspective will suggest further expanded cross-disciplinary tests of the ETH of disease. While the outcome of artificial selection for differential capacity for energy transfer is consistent with the ETH, explicit tests are needed.

## Acknowledgements

The LCR-HCR rat model system was supported by the Office of Research Infrastructure Programs/OD grant ROD012098A (to LGK and SLB) from the National Institutes of Health. Contact LGK or SLB for information on the LCR and HCR rats: these rat models are maintained as an international collaborative resource at the University of Toledo, Toledo, Ohio.

We are grateful to Peter Mitchell, Bruce Walsh, Jonathon Flint, Richard Mott, John Rapp, and James Watson for insights on the development of ideas presented here.

